# Learning inverse folding from millions of predicted structures

**DOI:** 10.1101/2022.04.10.487779

**Authors:** Chloe Hsu, Robert Verkuil, Jason Liu, Zeming Lin, Brian Hie, Tom Sercu, Adam Lerer, Alexander Rives

## Abstract

We consider the problem of predicting a protein sequence from its backbone atom coordinates. Machine learning approaches to this problem to date have been limited by the number of available experimentally determined protein structures. We augment training data by nearly three orders of magnitude by predicting structures for 12M protein sequences using AlphaFold2. Trained with this additional data, a sequence-to-sequence transformer with invariant geometric input processing layers achieves 51% native sequence recovery on structurally held-out backbones with 72% recovery for buried residues, an overall improvement of almost 10 percentage points over existing methods. The model generalizes to a variety of more complex tasks including design of protein complexes, partially masked structures, binding interfaces, and multiple states.

## 1. Introduction

Designing novel amino acid sequences that encode proteins with desired properties, known as *de novo protein design*, is a central challenge in bioengineering (Huang et al., 2016). The most well-established approaches to this problem use an energy function which directly models the physical basis of a protein’s folded state (Alford et al., 2017).

Recently a new class of deep learning based approaches has been proposed, using generative models to predict sequences for structures (Ingraham et al., 2019; Strokach et al., 2020; Anand-Achim et al., 2021; Jing et al., 2021b), generate backbone structures (Anand & Huang, 2018; Eguchi et al., 2020), jointly generate structures and sequences (Anishchenko et al., 2021; Wang et al., 2021), or model sequences directly (Rives et al., 2021; Madani et al., 2021; Shin et al., 2021; Gligorijevic et al., 2021; Bryant et al., 2021; Dallago et al., 2021). The potential to learn the rules of protein design directly from data makes deep generative models a promising alternative to current physics-based energy functions.

However, the relatively small number of experimentally determined protein structures places a limit on deep learning approaches. Experimentally determined structures cover less than 0.1% of the known space of protein sequences. While the UniRef sequence database (Suzek et al., 2015) has over 50 million clusters at 50% sequence identity; as of January 2022, the Protein Data Bank (PDB) (Berman et al., 2000) contains structures for fewer than 53,000 unique sequences clustered at the same level of identity.

Here we explore whether predicted structures can be used to overcome the limitation of experimental data. With progress in protein structure prediction (Jumper et al., 2021; Evans et al., 2022; Baek et al., 2021), it is now possible to consider learning from predicted structures at scale. Predicting structures for the sequences in large databases can expand the structural coverage of protein sequences by orders of magnitude. To train an inverse model for protein design, we predict structures for 12 million sequences in UniRef50 using AlphaFold2.

We focus on the problem of predicting sequences from backbone structures, known as *inverse folding* or fixed backbone design. We approach inverse folding as a sequence-to-sequence problem (Ingraham et al., 2019), using an autoregressive encoder-decoder architecture, where the model is tasked with recovering the native sequence of a protein from the coordinates of its backbone atoms.

We make use of the large number of sequences with unknown structures by adding them as additional training data, conditioning the model on predicted structures when the experimental structures are unknown (Figure 1). This approach parallels back-translation (Sennrich et al., 2015; Edunov et al., 2018) in machine translation, where predicted translations in one direction are used to improve a model in the opposite direction. Back-translation has been found to effectively learn from extra target data (i.e. sequences) even when the predicted inputs (i.e. structures) are of low quality.

**Figure 1.**
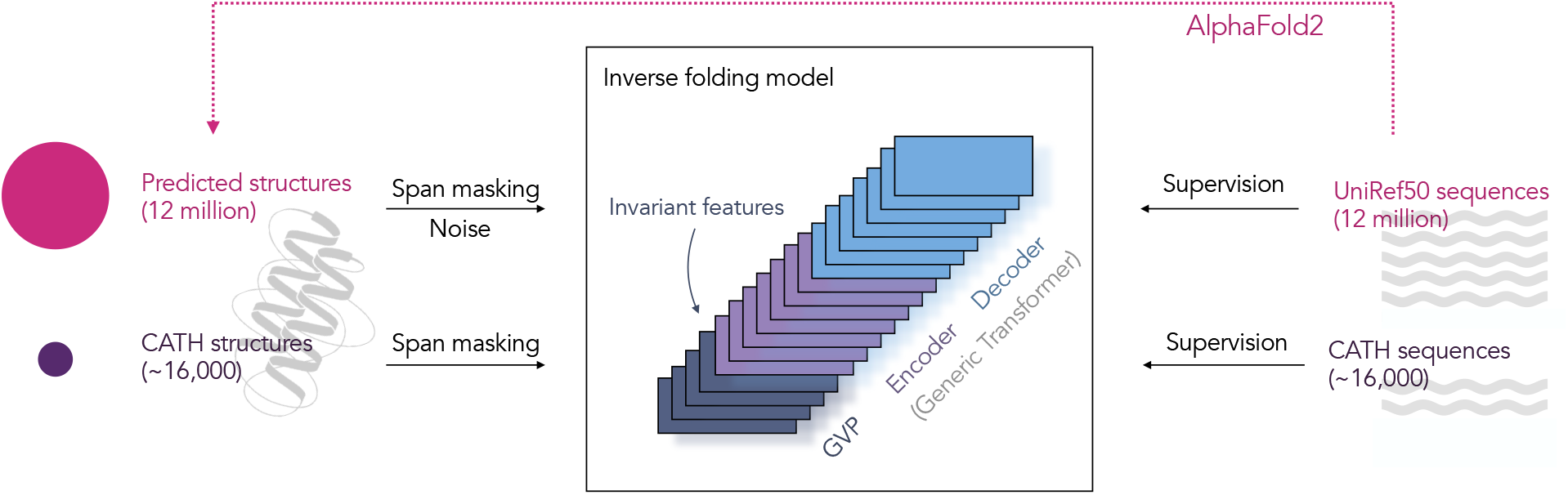
Augmenting inverse folding with predicted structures. To evaluate the potential for training protein design models with predicted structures, we predict structures for 12 million UniRef50 protein sequences using AlphaFold2 (Jumper et al., 2021). An autoregressive inverse folding model is trained to perform fixed-backbone protein sequence design. Train and test sets are partitioned at the topology level, so that the model is evaluated on structurally held-out backbones. We compare transformer models having invariant geometric input processing layers, with fully geometric models used in prior work. Span masking and noise is applied to the input coordinates.

**Figure 2.**
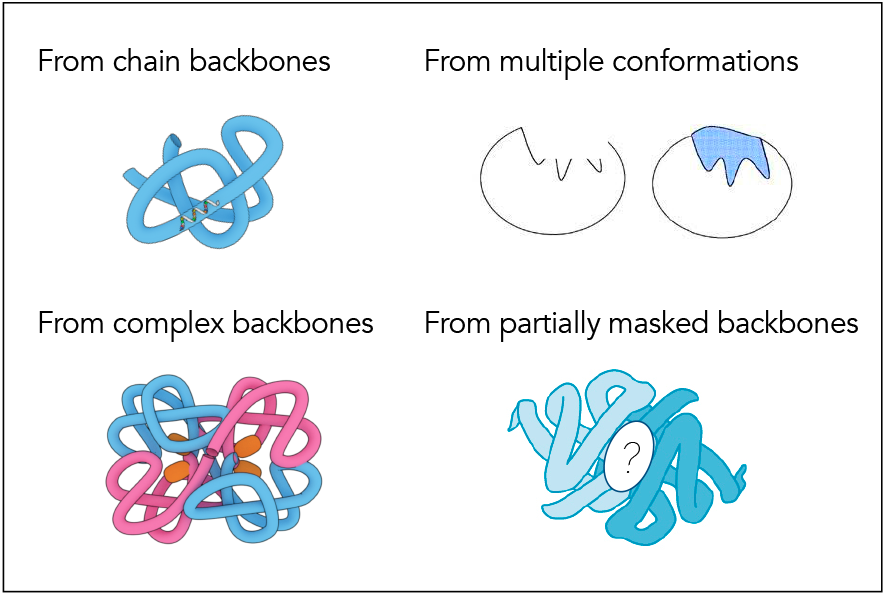
Illustration of the protein design tasks considered.

We find that existing approaches have been limited by data. While current state-of-the-art inverse folding models degrade when training is augmented with predicted structures, much larger models and different model architectures can effectively learn from the additional data, leading to an improvement of nearly 10 percentage points in the recovery of sequences for structurally held out native backbones.

We evaluate models on fixed backbone design benchmarks from prior work, and assess the generalization capabilities across a series of tasks including design of complexes and binding sites, partially masked backbones, and multiple conformations. We further consider the use of the models for zero-shot prediction of mutational effects on protein function and stability, complex stability, and binding affinity.

### 2. Learning inverse folding from predicted structures

The goal of inverse folding is to design sequences that fold to a desired structure. In this work, we focus on the backbone structure without considering side chains. While each of the 20 amino acid has a specific side chain, they share a common set of atoms that make up the amino acid backbone. Among the backbone atoms, we choose the N, C*α* (alpha Carbon), and C atom coordinates to represent the backbone.

Using the structures of naturally existing proteins we can train a model for this task by supervising it to predict the protein’s native sequence from the coordinates of its backbone atoms in three-dimensional space. Formally we represent this problem as one of learning the conditional distribution *p*(*Y|X*), where for a protein of length *n*, given a sequence *X* of spatial coordinates (*x*_1_, …, *x_i_*, …, *x*_3*n*_) for each of the backbone atoms *N, Cα, C* in the structure, the objective is to predict *Y* the native sequence (*y*_1_, …, *y_i_*, …, *y_n_*) of amino acids. This density is modeled autoregressively through a sequence-to-sequence encoder-decoder:

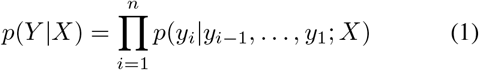

We train a model by minimizing the negative log likelihood of the data. We can design sequences by sampling, or by finding sequences that maximize the conditional probability given the desired structure.

### 2.1. Data

#### Predicted structures

We generate 12 million structures for sequences in UniRef50 to explore how predicted structures can improve inverse folding models. To select sequences for structure prediction we first use MSA Transformer (Rao et al., 2021) to predict distograms for MSAs of all UniRef50 sequences. We rank the sequences by distogram LDDT scores (Senior et al., 2020) as a proxy for the quality of the predictions. We take the top 12 million sequences not longer than five hundred amino acids and forward fold them using the AlphaFold2 model with a final Amber (Hornak et al., 2006) relaxation. This results in a predicted dataset approximately 750 times the size of the training set of experimental structures (Appendix A.1).

#### Training and evaluation data

We evaluate models on a structurally held-out subset of CATH (Orengo et al., 1997). We partition CATH at the topology level with an 80/10/10 split resulting in 16153 structures assigned to the training set, 1457 to the validation set, and 1797 to the test set. Particular care is required to prevent leakage of information in the test set via the predicted structures. We use Gene3D topology classification (Lees et al., 2012) to filter both the sequences used for supervision in training, as well as the MSAs used as inputs for AlphaFold2 predictions (Appendix A.1). We also perform evaluations on a smaller subset of the CATH test set that has been additionally filtered by TM-score using Foldseek (van Kempen et al., 2022) to exclude any structures with similarity to those in the training set (Appendix B).

### 2.2. Model architectures

We study model architectures using Geometric Vector Perceptron (GVP) layers (Jing et al., 2021b) that learn rotation-equivariant transformations of vector features and rotation-invariant transformations of scalar features.

We present results for three model architectures: (1) GVP-GNN from Jing et al. (2021b) which is currently state-of-the-art on inverse folding; (2) a GVP-GNN with increased width and depth (GVP-GNN-large); and (3) a hybrid model consisting of a GVP-GNN structural encoder followed by a generic transformer (GVP-Transformer). All models used in evaluations are trained to convergence, with detailed hyperparameters listed in Table A.1.

In inverse folding, the predicted sequence should be independent of the reference frame of the structural coordinates. For any rotation and translation *T* of the input coordinates, we would like for the model’s output to be invariant under these transformations, i.e., *p*(*Y|X*) = *p*(*Y|TX*). Both the GVP-GNN and GVP-Transformer inverse folding models studied in this work are invariant (Appendix A.3).

#### GVP-GNN

We start with the GVP-GNN architecture with 3 encoder layers and 3 decoder layers as described in (Jing et al., 2021b), with the vector gates described in (Jing et al., 2021a) (GVP-GNN, 1M parameters). As inputs to GVP-GNN, protein structures are represented as proximity graphs where each amino acid corresponds to a node in the graph. The node features are a combination of scalar node features derived from dihedral angles and vector node features derived from the relative positions of the backbone atoms, while the edge features capture the relative positions of nearby amino acids.

When trained on predicted structures, we find a deeper and wider version of GVP-GNN with 8 encoder layers and 8 decoder layers (GVP-GNN-large, 21M parameters) performs better. Scaling GVP-GNN further did not improve model performance in preliminary experiments (Figure 6c).

#### GVP-Transformer

We use GVP-GNN encoder layers to extract geometric features, followed by a generic autoregressive encoder-decoder Transformer (Vaswani et al., 2017). In GVP-GNN, the input features are translation-invariant and each layer is rotation-equivariant. We perform a change of basis on the vector features from GVP-GNN into local reference frames defined for each amino acid to derive rotation-invariant features (Appendix A.3). In ablation studies increasing the number of GVP-GNN encoder layers improves the overall model performance (Figure C.1), indicating that the geometric reasoning capability in GVP-GNN is complementary to the Transformer layers. Scaling improves performance up to a 142M-parameter GVP-Transformer model with 4 GVP-GNN encoder layers, 8 generic Transformer encoder layers, and 8 generic Transformer decoder layers (Figure 6c).

### 2.3. Training

#### Combining experimental and predicted data

During training, in each epoch we mix the training set of experimentally derived structures (~ 16K structures) with a 10% random sample of the AlphaFold2-predicted training set (10% of 12M), resulting in a 1:80 experimental:predicted data ratio. For larger models, a high ratio of predicted data during training helps prevent overfitting on the smaller experimental train set (Figure 6b). While adding predicted data improves performance, training only on predicted data leads to substantially worse performance (Table C.2).

The loss is equally weighted for each amino acid in target sequences. We mask out predicted input coordinates with AlphaFold2 confidence score (pLDDT) below 90, around 25% of the predicted coordinates. See Figure 3 for visualization of the pLDDT confidence score. Most often these low confidence regions are at the start and the end of sequences and may correspond to disordered regions. We prepend one token at the beginning of each sequence to indicate whether the structure is experimental or predicted. For each residue we provide the pLDDT confidence score from AlphaFold2 as a feature encoded by Gaussian radial basis functions.

**Figure 3.**
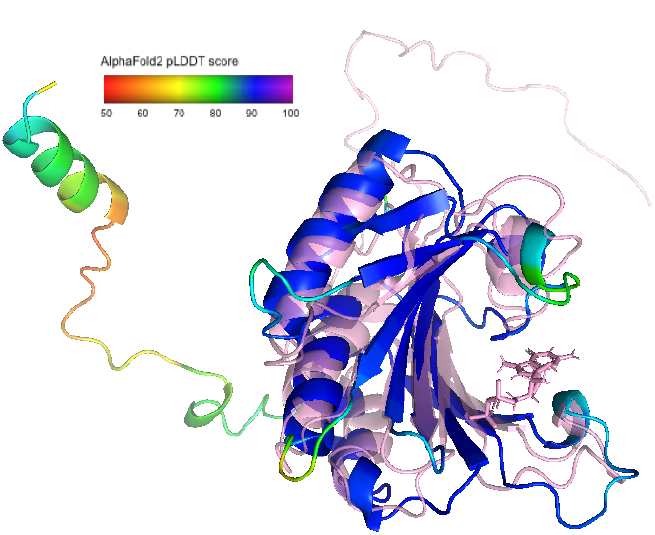
Example AlphaFold prediction compared with experimental structure for a UniRef50 sequence (UniRef50: P07260; PDB: 1AP8). The experimental structure is shown as pink with transparency. The prediction is coloured by the pLDDT confidence score, with blue in high-confidence regions.

Adding Gaussian noise at the scale of 0.1 angstroms to the predicted structures during training slightly improves performance (Table C.1). This finding is consistent with Edunov et al. (2018), who observe that backtranslation with sampled or noisy synthetic data provides a stronger training signal than maximum a posteriori (MAP) predictions.

#### Span masking

To enable sequence design for partially masked backbones, we introduce backbone masking during training. We experiment with both independent random masking and span masking. In natural language processing, span masking improves performance over random masking (Joshi et al., 2020). We randomly select continuous spans of up to 30 amino acids until 15% of input backbone coordinates are masked. The communication patterns in the geometric layers are adapted to account for masking with details in Appendix A.2. Span masking improves the performance of GVP-Transformer both on unmasked backbones (Table C.1) and on masked regions (Figure 4).

**Figure 4.**
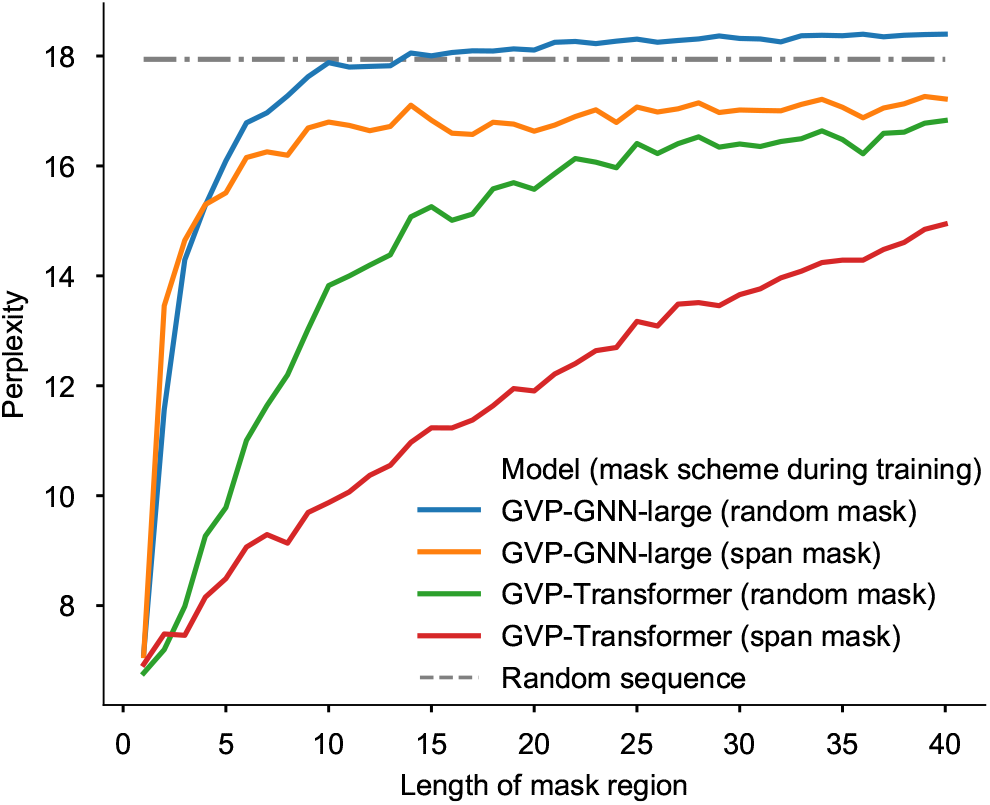
Perplexity on regions of masked coordinates of different lengths. The GVP-GNN architecture degrades to the perplexity of the background distribution for masked regions of more than a few tokens, while GVP-Transformer maintains moderate accuracy on long masked spans, especially when trained on masked spans.

## 3. Results

We evaluate models across a variety of benchmarks in two overall settings: fixed backbone sequence design and zero-shot prediction of mutation effects. For fixed backbone design, we start with evaluation in the standard setting (Ingraham et al., 2019; Jing et al., 2021b) of sequence design given all backbone coordinates. Then, we make the sequence design task more challenging along three dimensions: (1) introducing masking on coordinates; (2) generalization to protein complexes; and (3) conditioning on multiple conformations. Additionally, we show that inverse folding models are effective zero-shot predictors for protein complex stability, binding affinity, and insertion effects.

### 3.1. Fixed backbone protein design

We begin with the task of predicting the native protein sequence given its backbone atom (N, C*α*, C) coordinates. Perplexity and sequence recovery on held-out native sequences are two commonly used metrics for this task. Perplexity measures the inverse likelihood of native sequences in the predicted sequence distribution (low perplexity for high likelihood). Sequence recovery (accuracy) measures how often sampled sequences match the native sequence at each position. To maximize sequence recovery, the predicted sequences are sampled with low temperature *T* = 1e − 6 from the model. While the model is calibrated (Figure C.5), a lower temperature results in sequences with higher likelihoods (and hence typically higher sequence recovery) and lower diversity. Empirically at temperature as low as 1e − 6 the sampling is almost deterministic. Table 1 compares models using the perplexity and sequence recovery metrics on the structurally held-out backbones.

**Table 1.** Fixed backbone sequence design. Evaluation on the CATH 4.3 topology split test set. Models are compared on the basis of per-residue perplexity (lower is better; lowest perplexity bolded) and sequence recovery (higher is better; highest sequence recovery bolded). Large models can make better use of the predicted UniRef50 structures. The best model trained with predicted structures (GVP-Transformer) improves sequence recovery by 8.9 percentage points over the best model (GVP-GNN) trained on CATH only.

| Model | Data | Perplexity |  |  | Recovery % |  |  |
| --- | --- | --- | --- | --- | --- | --- | --- |
|  |  | Short | Single-chain | All | Short | Single-chain | All |
| Natural frequencies |  | 18.12 | 18.03 | 17.97 | 9.6% | 9.0% | 9.5% |
| Structured GNN | CATH | 7.91 | 6.48 | 6.49 | 31.5% | 37.1% | 37.1% |
| GVP-GNN | CATH | 7.14 | 5.36 | 5.43 | 34.0% | 42.7% | 42.2% |
|  | + AlphaFold2 | 8.55 | 6.17 | 6.06 | 29.5% | 38.2% | 38.6% |
| GVP-GNN-large | CATH | 7.68 | 6.12 | 6.17 | 32.6% | 39.4% | 39.2% |
|  | + AlphaFold2 | 6.11 | 4.09 | 4.08 | **38.3%** | 50.8% | 50.8% |
| GVP-Transformer | CATH | 8.18 | 6.33 | 6.44 | 31.3% | 38.5% | 38.3% |
|  | + AlphaFold2 | **6.05** | **4.00** | **4.01** | 38.1% | **51.5%** | **51.6%** |

We observe that current state-of-the-art inverse folding models are limited by the CATH training set. Scaling the current 1M parameter model (GVP-GNN) to 21M parameters (GVP-GNN-large) on the CATH dataset results in overfitting with a degradation of sequence recovery from 42.2% to 39.2% (Table 1). On the other hand, the current model at the 1M parameter scale cannot make use of the predicted structures: training GVP-GNN with predicted structures results in a degradation to 38.6% sequence recovery (Table 1), with performance worsening with increasing numbers of predicted structures in training (Figure 6a).

Larger models benefit from training on the AlphaFold2-predicted UniRef50 structures. Training with predicted structures increases sequence recovery from 39.2% to 50.8% for GVP-GNN-large and from 38.3% to 51.6% for GVP-Transformer over training only on the experimentally derived structures. The improvements are also reflected in perplexity. Similar improvements are observed on the test subset filtered by TM-score (Table B.1). The best model trained with UniRef50 predicted stuctures, GVP-Transformer, improves sequence recovery by 9.4 percentage points over the best model, GVP-GNN, trained on CATH alone.

As there are many sequences that can fold to approximately the same structure, even an ideal protein design model will not have 100% native sequence recovery. We observe that the GVP-GNN-large and GVP-Transformer models are well-calibrated (Figure C.5). The substitution matrix between native sequences and model-designed sequences resembles the BLOSUM62 substitution matrix (Figure C.4), albeit noticeably sparser for the amino acid Proline.

When we break down performance on core residues and surface residues, as expected, core residues are more constrained and have a high native sequence recovery rate of 72%, while surface residues are not as constrained and have a lower sequence recovery of 39% (Figure 5; top). Generally perplexity increases with the solvent accessible surface area (Figure 5; bottom). Despite the lower sequence recovery on the surface, sampled sequences do tend not to have hydrophobic residues on the surface (Figure C.6).

**Figure 5.**
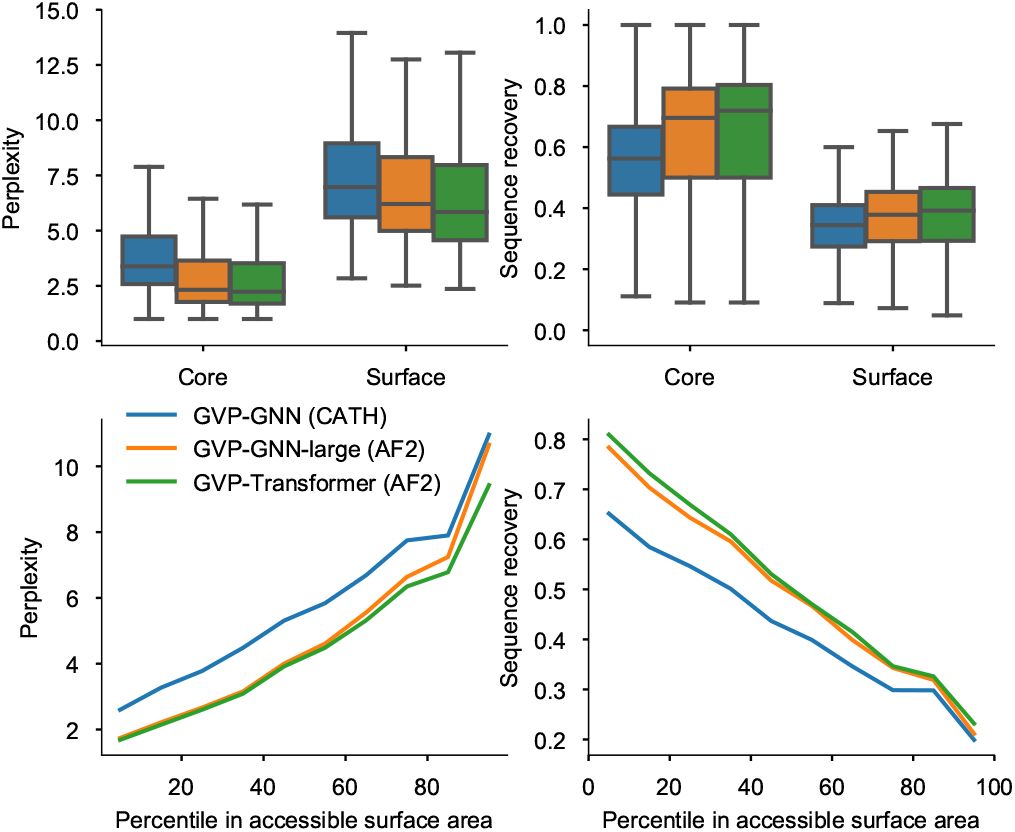
Comparison of perplexity and sequence recovery by structural context according to two different measures: number of neighbors (top) and solvent accessible surface area (bottom). Top: Breakdown for core and surface residues. Residues are categorized by density of neighboring C*α* atoms within 10A of the central residue C*α* atom (core: ≥ 24 neighbors; surface: < 16 neighbors). Each box shows the distribution of perplexities for the core or surface residues across different sequences. Bottom: Perplexity and sequence recovery as a function of solvent accessible surface area. Increased sequence recovery for buried residues suggests the model learns dense hydrophobic packing constraints in the core.

**Figure 6.**
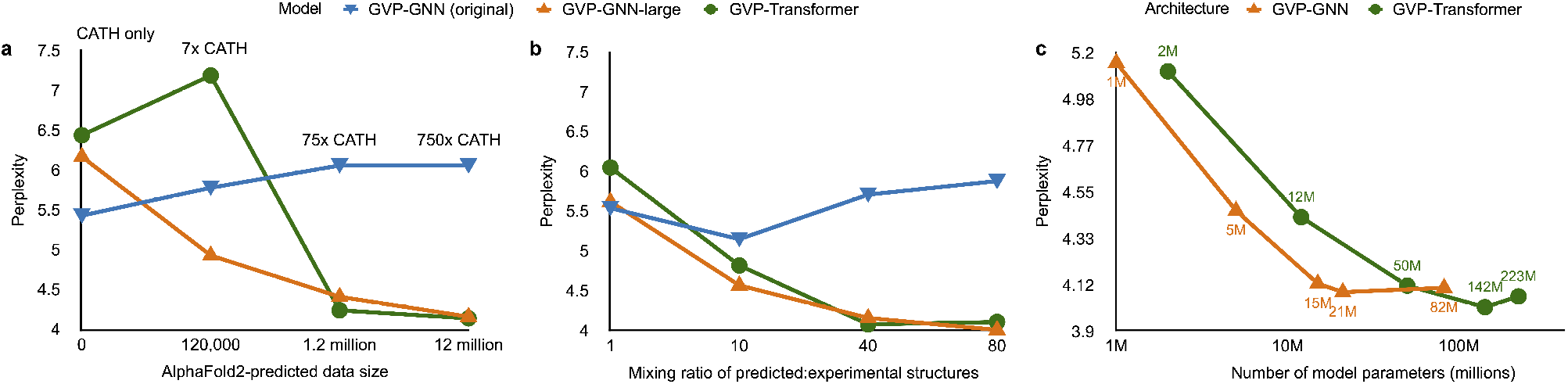
Ablation studies on training data. (a) Effect of increasing the number of predicted structures. The original GVP-GNN degrades with training on additional data, but GVP-GNN-large and GVP-Transformer improve with increasing numbers of predicted structures. (b) Effect of increasing the mixing ratio during training between predicted and experimental structures. A higher ratio of predicted structures improves performance for both GVP-GNN-large and GVP-Transformer. (c) GVP-GNN and GVP-Transformer model size.

As an example of inverse folding of a structurally-remote protein, we re-design the receptor binding domain (RBD) sequence of the SARS-CoV-2 spike protein (PDB: 6XRA and 6VXX; illustrated in Figure C.3) with the two models. The SARS-CoV-2 spike protein has no match to the training data with TM-score above 0.5. Both GVP-GNN and GVP-Transformer achieve high sequence recovery (49.7% and 53.6%) for the native RBD sequence (Table C.4). See Table C.8 for a list of randomly sampled sequence designs.

While perplexity and sequence recovery are informative metrics, low perplexity and high sequence recovery do not necessarily guarantee structural similarity. One empirically observed failure mode in sampled sequences is repetition of the same amino acid, e.g. EEEEEEE. It would be interesting to further identify more failure modes by studying the experimental or predicted structures of sampled sequences.

#### Partially-masked backbones

We evaluate the models on partial backbones. While masking during training does not significantly change test performance on unmasked backbones (Table C.1), masking does enable models to non-trivially predict sequences for mask regions. Although GVP-GNN-large has low perplexity on short-length masks, its performance quickly degrades to the perplexity of the background distribution on masks longer than 5 amino acids (Figure 4). By contrast, the GVP-Transformer model maintains moderate performance even on longer masked regions, with less degradation if trained with span masking instead of independent random masking (Figure 4).

#### Protein complexes

Although the training data only consists of single chains, we find that models generalize to multi-chain protein complexes. We represent complexes by concatenating the chains together with 10 mask tokens between chains, and place the target chain for sequence design at the beginning during concatenation. We include all complexes in the CATH 4.3 test set up to 1000 amino acids in length. For chains that are part of a protein complex, there is a substantial improvement in perplexity of both models when given the full complex coordinates as input, versus only the single chain (Table 2 and Figure C.2), suggesting that both GVP-GNN and GVP-Transformer can make use of inter-chain information from amino acids that are close in 3D structure but far apart in sequence.

**Table 2.**
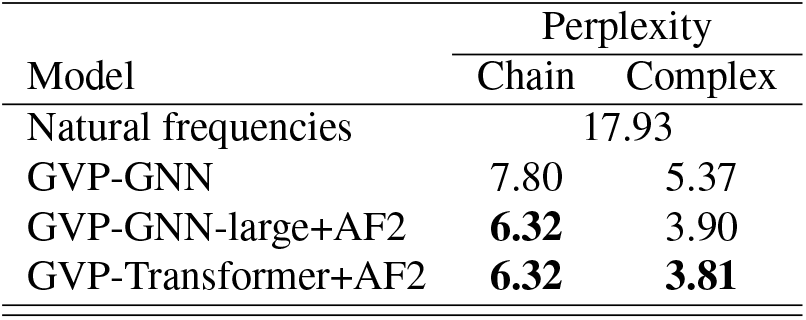
Sequence design performance on complexes in the CATH topology test split when given the backbone coordinates of only a chain (“Chain” column) and when given all backbone coordinates of the complex (“Complex” column). The perplexity is evaluated on the same chain in the complex for both columns.

#### Multiple conformation

Multi-state design is of interest for engineering enzymes and biosensors (Langan et al., 2019; Quijano-Rubio et al., 2021). Some proteins exist in multiple distinct folded forms in equilibrium, while other proteins may exhibit distinct conformations when binding to partner molecules.

For a backbone *X*, the inverse folding model predicts a conditional distribution *p*(*Y |X*) over possible sequences *Y* for the backbone. To design a protein sequence compatible with two states *A* and *B*, we would like find sequences with high likelihoods in the conditional distributions *p*(*Y |A*) and *p*(*Y |B*) for each state. We use the geometric average of the two conditional likelihoods as a proxy for the desired distribution *p*(*Y|A, B*) conditioned on the sequence being compatible with both states.

We compare single-state and multi-state sequence design performance on 87 test split proteins with multiple conformations in the PDBFlex dataset (Hrabe et al., 2016). On locally flexible residues, multi-state design results in lower sequence perplexity than single-state design (Figure 7). See Appendix C for more details on the PDBFlex data.

**Figure 7.**
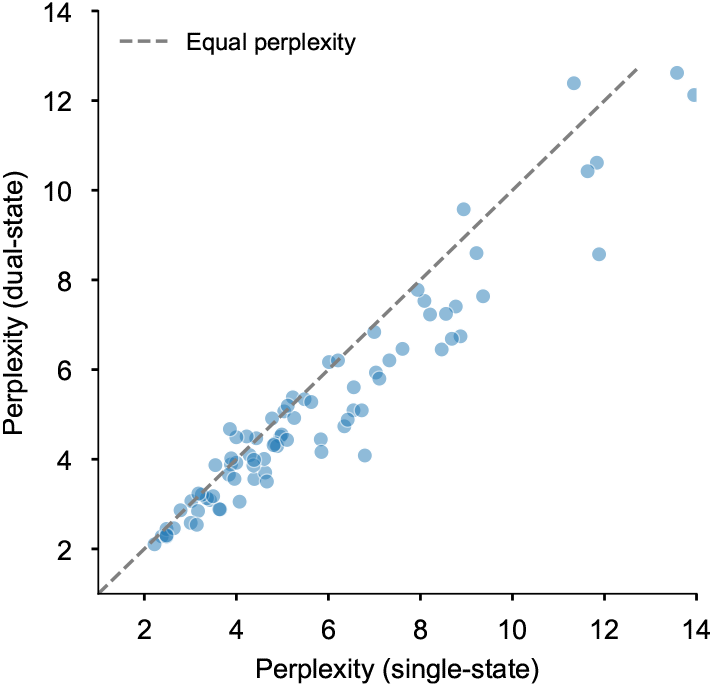
Dual-state design. GVP-Transformer conditioned on two conformations results in lower sequence perplexity at locally flexible residues than single-conformation conditioning for structurally held-out proteins in PDBFlex (see Appendix C for details).

### 3.2. Zero-shot predictions

We next show that inverse folding models are effective zero-shot predictors of mutational effects across practical design applications, including prediction of complex stability, binding affinity, and insertion effects. To score the effect of a mutation on a particular sequence, we use the ratio between likelihoods of the mutated and wildtype sequences according to the inverse folding model, given the experimentally determined wildtype structure. Exact likelihood evaluations are possible from both GVP-GNN and GVP-Transformer as they are both based on autoregressive decoders. We then compare these likelihood ratio scores to experimentally-determined fitness values measured on the same set of sequences.

#### *De novo* mini-proteins

Rocklin et al. (2017) performed deep mutational scans across a set of *de novo* designed mini-proteins with 10 different folds measuring the stability in response to point mutations. The likelihoods of inverse folding models have been shown to correlate with experimentally measured stability using this dataset (Ingraham et al., 2019; Jing et al., 2021b). We evaluate the GVP-Transformer and GVP-GNN-large models on the same mutational scans, and observe improvements in stability predictions from using predicted structures as training data for 8 out of 10 folds in the dataset (Table C.3). Further details are in Appendix C.

#### Complex stability

We evaluate models on zero-shot prediction of mutational effects on protein complex interfaces, using the Atom3D benchmark (Townshend et al., 2020) which incorporates binding free energy changes in the SKEMPI database (Jankauskaitė et al., 2019) as a binary classification task. We find that sequence log-likelihoods from GVP-GNN are effective zero-shot predictors of stability changes of protein complexes even without predicted structures as training data (Table C.5), performing comparably to the best supervised method which uses transfer learning. While we observe a substantial improvement in perplexity when predicted structures are added to training (Table 2), this does not further improve complex stability prediction for the single-point mutations in SKEMPI (Table C.5), indicating potential limitations of evaluating models only on single-point mutations.

#### Binding affinity

While the SKEMPI dataset features one mutation entry per protein, we also want to evaluate whether inverse folding models can rank different mutations on the same protein, potentially enabling binding-affinity optimization, which is an important task in therapeutic design. We assess whether inverse folding models can predict mutational effects on binding by leveraging a dataset generated by Starr et al. (2020) in which all single amino acid substitutions to the SARS-CoV-2 receptor binding domain (RBD) were experimentally measured for binding affinity to human ACE2. Given potential applications to interface optimization or design, we focus on mutations within the receptor binding motif (RBM), the portion of the RBD in direct contact with ACE2 (Lan et al., 2020). When given all RBD and ACE2 coordinates, the best inverse folding model produces RBD-sequence log-likelihoods that have a Spearman correlation of 0.69 with experimental binding affinity measurements (Table 3). We observe weaker correlations when not providing the model with ACE2 coordinates, indicating that inverse folding models take advantage of structural information in the binding partner. When masking RBM coordinates (69 of 195 residues, a longer span than masked during model training), we no longer observe correlation between RBD log-likelihood and binding affinity, indicating that the model relies on structural information at the interface to identify interface designs that preserve binding. Zero-shot prediction via inverse folding outperforms methods for sequence-based variant effect prediction, which use the likelihood ratio between the mutant and wildtype amino acids at each position to predict the impact of a mutation on binding affinity. These likelihoods are inferred by masked language models, ESM-1b, ESM-1v, and ESM-MSA-1b, as described by Meier et al. (2021) (Table 3); additional details are given in Appendix C.

**Table 3.** Zero-shot performance on binding affinity prediction for the receptor binding domain (RBD) of SARS-CoV-2 Spike, evaluated on ACE2-RBD mutational scan data (Starr et al., 2020). The zero-shot predictions are based on the sequence log-likelihood for the receptor binding motif (RBM), which is the portion of the RBD in direct contact with ACE2 (Lan et al., 2020). We evaluate in four settings: 1) Given sequence data alone (“No coords”); 2) Given backbone coordinates for both ACE2 and the RBD but excluding the RBM and without sequence (“No RBM coords”); 3) Given the full backbone for the RBD but no information for ACE2 (“No ACE2 coords”); and 4) Given all coordinates for the RBD and ACE2.

| Model | Spearman correlation |  |  |  |
| --- | --- | --- | --- | --- |
|  | No coords | No RBM coords | No ACE2 coords | All coords |
| ESM-1v | 0.03 |  |  |  |
| ESM-1b | 0.02 |  |  |  |
| ESM-MSA-1b (few-shot) | **0.51** |  |  |  |
| GVP-GNN |  | -0.10 | 0.50 | 0.60 |
| GVP-GNN-large+AF2 |  | -0.05 | 0.52 | **0.69** |
| GVP-Transformer+AF2 |  | -0.06 | **0.53** | 0.64 |

#### Sequence insertions

Using masked coordinate tokens at insertion regions, inverse folding models can also predict insertion effects. On adeno-associated virus (AAV) capsid variants, we show that relative differences in sequence log-likelihoods correlate with the experimentally measured insertion effects from Bryant et al. (2021). As shown in Table C.6, both GVP-GNN and GVP-Transformer outperform the sequence-only zero-shot prediction baseline ESM-1v (Meier et al., 2021). When evaluating on subsets of s quences increasingly further away from the wildtype (≥ 2, ≥ 3, and ≥ 8 mutations), the GVP-GNN-large and GVP-Transformer models trained with predicted structures have increasing advantages compared to GVP-GNN trained without predicted structures.

## 4. Related work

### Structure-based protein sequence design

Early work on design of protein sequences studied the packing of amino acid side chains to fill the interior space of predetermined backbone structures, either for a fixed backbone conformation (Street & Mayo, 1999; Dahiyat & Mayo, 1997; DeGrado et al., 1991), or with flexibility in the backbone conformation (Harbury et al., 1998). Since then, the Rosetta energy function (Alford et al., 2017) has become an established approach for structure-based sequence design. An alternative non-parametric approach involves decomposing the library of known structures into common sequence-structure motifs (Zhou et al., 2020).

Early machine learning approaches in structure-based protein sequence design used fragment-based and energy-based global features derived from structures (Li et al., 2014; O’Connell et al., 2018). More recently, convolution-based deep learning methods have also been applied to predict amino acid propensities given the surrounding local structural environments (Anand-Achim et al., 2021; Boomsma & Frellsen, 2017; Shroff et al., 2020; Li et al., 2020; Qi & Zhang, 2020; Zhang et al., 2020; Chen et al., 2019; Wang et al., 2018). Another recent machine learning approach is to leverage structure prediction networks for sequence design. Anishchenko et al. (2021) carried out Monte Carlo sampling in the sequence space to invert the trRosetta (Yang et al., 2020) structure prediction network for sequence design, while Norn et al. (2021) backpropagated gradients through the trRosetta network.

#### Generative models of proteins

The literature on structure-based generative models of protein sequences is the closest to our work. Ingraham et al. (2019) introduced the formulation of fixed-backbone design as a conditional sequence generation problem, using invariant features with graph neural networks, modeling each amino acid as a node in the graph with edges connecting spatially adjacent amino acids. Jing et al. (2021b;a) further improved graph neural networks for this task by developing architectures with translation- and rotation-equivariance to enable geometric reasoning, showing that GVP-GNN achieves higher native sequence recovery rates than Rosetta on TS50, a benchmark set of 50 protein chains. Strokach et al. (2020) trained graph neural networks for conditional generation with the masked language modeling objective, adding homologous sequences as data augmentation to training. Most recently, contemporary with our work, Dauparas et al. (2022) improved an existing graph neural network (Ingraham et al., 2019) by combining additional input features and edge updates, and validated designed protein sequences through X-ray crystallography, cryoEM and functional studies. Also contemporary with our work, Yang et al. (2022) showed that pretrained sequence-only protein masked language models can be combined with structure-based graph neural networks to improve inverse folding performance.

Recently models have been proposed to jointly generate structures and sequences. Anishchenko et al. (2021) generate structures by optimizing sequences through the trRosetta structure prediction network to maximize their difference from a background distribution. The joint generation approach is also being explored in the setting of infilling partial structures. Contemporary to this work, Wang et al. (2021) apply span masking to fine-tune the RosettaFold model (Baek et al., 2021) to perform infilling, although conditioning on both coordinates and amino acid identities instead of considering the inverse folding task. Also contemporary to this work, Jin et al. (2021) develop a conditional generation model for jointly generating sequences and structures for antibody complementarity determining regions (CDRs), conditioned on framework region structures. Anand & Achim (2022) introduced an equivariant denoising diffusion approach for jointly generating protein structures and sequences for infilling, loop completion, and beyond.

So far there has been little work on generative models of structures directly. Interesting examples include Anand & Huang (2018) who model fixed-length protein backbones with generative adversarial networks (GANs) via pairwise distance matrices, and Eguchi et al. (2020) who generate antibody structures with variational autoencoders (VAEs).

#### Language models

A large body of work has focused on modeling the sequences in individual protein families. Shin et al. (2021) show that protein-specific autoregressive sequence models trained on related proteins can predict point mutation and indel effects and design functional nanobodies. Trinquier et al. (2021) also studied protein-specific autoregressive models for sequence generation.

Recently language models have been proposed for modeling large scale databases of protein sequences rather than families of related sequences. Examples include (Bepler & Berger, 2019; Alley et al., 2019; Heinzinger et al., 2019; Rao et al., 2019; Madani et al., 2020; Elnaggar et al., 2021; Rives et al., 2021; Rao et al., 2021). Meier et al. (2021) found that the log-likelihoods of large protein language models predict mutational effects. Madani et al. (2021) study an autoregressive sequence model conditioned on functional annotations and show it can generate functional proteins.

#### Structure-agnostic protein sequence design

We point the reader to Wu et al. (2021) for a review of the many machine learning-based sequence design approaches that do not explicitly model protein structures. Additionally, as an alternative to sequence generation models, model-guided algorithms design sequences based on predictive models as oracles (Yang et al., 2019; Angermueller et al., 2019; Brookes et al., 2019; Sinai et al., 2020).

#### Back-translation

For machine translation (MT) in NLP, Sennrich et al. (2015) studied how to leverage large amounts of monolingual data in the target language, a setting that parallels the situation we consider with protein sequences (the target language in our case). Sennrich et al. found it most effective to generate synthetic source sentences by performing the backwards translation from the target sentence, i.e. back-translation. This parallels the approach we take of predicting structures for sequence targets that have unknown structures. Edunov et al. (2018) further investigated back-translation for large-scale language models.

### 5. Conclusions

While there are billions of protein sequences in the largest sequence databases, the number of available experimentally determined structures is on the order of hundreds of thousands, imposing a limit on generative methods that learn from protein structure data. In this work, we explored whether predicted structures from recent deep learning methods can be used in tandem with experimental structures to train models for protein design.

To this end, we generated structures for 12 million UniRef50 sequences using AlphaFold2. As a result of training with this data we observe improvements in perplexity and sequence recovery by substantial margins, and demonstrate generalization to longer protein complexes, to proteins in multiple conformations, and to zero-shot prediction for mutation effects on binding affinity and AAV packaging. These results highlight that in addition to the geometric inductive biases which have been the major focus for work on inverse-folding to date, finding ways to leverage more sources of training data is an equally important path to improved modeling capabilities.

Contemporary with our work, the AlphaFold Protein Structure Database (Varadi et al., 2021) is rapidly growing, featuring 1 million predicted structures as of June 2022. Structure-based protein design models will likely continue to benefit from this new data source as the coverage expands to encompass more of the structural universe. Additionally, with the recent progress in structure prediction for multi-chain protein complexes (Evans et al., 2022; Humphreys et al., 2021), predicted complex structures could be another valuable source of data for learning protein-protein interactions.

We also take initial steps toward more general structure-conditional protein design tasks. By integrating backbone span masking into the inverse folding task and using a sequence-to-sequence transformer, reasonable sequence predictions can be achieved for short masked spans.

If ways can be found to continue to leverage predicted structures for generative models of proteins, it may be possible to create models that learn to design proteins from an expanded universe of the billions of natural sequences whose structures are currently unknown.

## Acknowledgements

We thank Halil Akin, Sal Candido, David Ding, Ori Kabeli, Joshua Meier, Sergey Ovchinnikov, Prajit Ramachandran, Ammar Rizvi, Kathy Wei, Kevin Yang, Zhongkai Zhu, and the anonymous reviewers for feed-back on the manuscript and insightful conversations.

## A. Additional details on datasets, training procedures, and model architectures

### A.1. Details on dataset of predicted structures

We used training data from two sources: 1) experimental protein structures from the CATH 40% non-redundant chain set, and 2) AlphaFold2-predicted structures from UniRef50 sequences. To evaluate the generalization performance across different protein folds, we split the train, validation, and test data based on the CATH hierarchical classification of protein structures (Orengo et al., 1997) for both data sources. To achieve that a rigorous structural hold-out, we additionally use foldseek (van Kempen et al., 2022) for pairwise TMalign between the test set the train set.

#### CATH topology split

Following the structural split methodology in previous work (Ingraham et al., 2019; Jing et al., 2021b; Strokach et al., 2020), we randomly split the CATH v4.3 (latest version) topology classification codes into train, validation, and test sets at a 80/10/10 ratio. The CATH (Orengo et al., 1997) structural hierarchy, classifies domains in four levels: Class (C), Architecture (A), Topology/fold (T), and Homologous superfamily (H). The topology/fold (T) level roughly corresponds to the SCOP fold classification.

#### Experimental structures

We collected full chains up to length 500 for all domains in the CATH v4.3 40% sequence identity non-redundant set. The experimental structure data contained only stand-alone chains and no multichain complexes. As each chain may be classified with more than one topology codes, we further removed chains with topology codes spanning different splits, so that there is no overlap in topology codes between train, validation, and test. This results in 16,153 chains in the train split, 1457 chains in the validation split, and 1797 chains in the test split.

#### Predicted structures

We curated a new data set of AlphaFold2 (Jumper et al., 2021)-predicted structures for a selective subset of UniRef50 (202001) sequences. To prevent information leakage about the test set from the predicted structures, we proceeded in the following steps.

First, we annotated UniRef50 sequences with CATH classification according to the Gene3D (Lees et al., 2012) database, also used by Strokach (Strokach et al., 2020) for data curation. Gene3D represents each CATH classification code as a library of representative profile HMMs. We searched all HMMs associated with the validation and test splits against the UniRef50 sequences using default parameters in hmmsearch (Potter et al., 2018) and excluded all hits.

Additionally, as AlphaFold2 predictions use multiple sequence alignments (MSAs) as inputs, we also took precaution to avoid information leakage from sequences in the MSAs. We created a filtered version of UniRef100 by searching all the validation-split and test-split Gene3D HMMs against UniRef100 (202001) and excluding all hits. Then, we constructed our MSAs using hhblits (Steinegger et al., 2019) on this filtered version of UniRef100. While this filtering step was out of precaution, in retrospect it was perhaps unlikely for the MSA inputs to AlphaFold2 to leak information as the MSAs themselves were not seen during training. The filtering step may have negatively impacted the quality of the resulting predicted structures, although empirically only a very small percentage of MSA sequences were filtered out.

As AlphaFold2 predictions are computationally costly, our budget only allowed for predicting structures for a subset of the UniRef50 sequences. We ranked UniRef50 sequences based on the distogram lDDT score, based on distogram predictions from MSATransformer (Rao et al., 2021), as a proxy for the quality of predicted structures. While the original distogram lDDT score (Supplementary Equation 6 in (Senior et al., 2020)) is based on pairwise distances from native protein structures, in the absence of native structures we use the argmax of pairwise distances instead, effectively measuring the “sharpness” of distograms and prioritizing sharper predictions. In this order, using AlphaFold2 Model 1 on the filtered UniRef100 MSAs described above, we obtained predicted structures for the top 12 million UniRef50 sequences under length 500, roughly 750 times the CATH train set size.

We used the publicly released model weights from AlphaFold2 Model 1 for CASP14 as a single model, as opposed the 5-model ensemble in (Jumper et al., 2021), to cover more sequences with the same amount of computing resources. We curated the input MSAs from UniRef100 with hhblits, with an additional filtering step as described above. To reduce computational costs, compared to the standard AlphaFold2 protocol, we did not include the UniRef90 jackhmmer MSAs, or the MGnify and BFD metagenomics MSAs, nor the pdb70 templates. Other than a reduced inputs, we followed the default settings in AlphaFold2 open source code, using 3 recycling iterations and the default Amber relaxation protocol. Despite the reduced inputs, the resulting 12 million predicted structures still have high pLDDT scores from AlphaFold, with 75% of residues having pLDDT above 90 (highly confident).

**Table A.1.** Details on model hyperparameters and training.

|  | GVP-GNN | GVP-GNN-large | GVP-Transformer |
| --- | --- | --- | --- |
| GVP-GNN embedding dim (node) | (100, 16) | (256, 64) | (1024, 256) |
| GVP-GNN embedding dim (edge) | (32, 1) | (32, 1) | (32, 1) |
| Top K neighbors in GVP-GNN | 30 | 30 | 30 |
| GVP-GNN encoder layers | 3 | 8 | 4 |
| GVP-GNN decoder layers | 3 | 8 |  |
| Transformer embedding dim |  |  | 512 |
| Feedforward embedding dim |  |  | 2048 |
| Attention heads |  |  | 8 |
| Transformer encoder layers |  |  | 8 |
| Transformer decoder layers |  |  | 8 |
| **Total number of parameters** | **1M** | **21M** | **142M** |
| Batch size (tokens per GPU) | 3072 | 4096 | 4096 |
| GPUs | 1 | 32 | 32 |
| CATH:AF2 mixing ratio | 1:0 | 40:1 | 80:1 |
| Epochs until convergence | 84 | 368 | 178 |
| Train time per epoch (GPU hours) | 0.07 | 24 | 88 |
| **Total train time (GPU days)** | **0.2** | **368** | **653** |
| Optimizer | Adam | Adam | Adam |
| Learning rate schedule | Constant | Inverse square root | Inverse square root |
| Peak learning rate | 1.0E-03 | 1.0E-03 | 1.0E-03 |
| Initial learning rate |  | 1.0E-07 | 1.0E-07 |
| Warm-up updates |  | 5000 | 5000 |
| Gradient clipping | 4.0 | 1 |  |

We found that increasing the predicted data size to up to 1 million structures (75 times the CATH experimental data size) substantially improves model performance. Beyond 1 million structures, models still benefit from more data but with diminished marginal returns (Figure 6a).

#### Noise on AlphaFold2-predicted backbone coordinates

Even after Amber relaxation, the backbone coordinates predicted by AlphaFold2 contain artifacts in the sub-Angstrom scale that may give away amino acid identities. Without adding noise on predicted structures, there is a substantial gap between held-out set performance on predicted structures and on experimental structures. To prevent the model from learning non-generalizable AlphaFold2-specific rules, we added Gaussian noise at the 0.1A scale on predicted backbone coordinates. The Gaussian noise improves the invariant Transformer performance but not the GVP-GNN performance (Supplementary Figure C.1).

### A.2. Details on span masking

We add a binary feature indicating whether each coordinate is masked or not. In GVP-Transformer, we exclude the masked nodes in the GVP-GNN encoder layers, and then impute zeros when passing the GVP-GNN outputs into the main Transformer. Imputing zeros for missing vector features ensure the rotation- and translation-invariance of the model. In GVP-GNN, we impute zeros for the input vector features, and in the input graph connect the masked nodes to the *k* sequence nearest-neighbors (*k* = 30) in lieu of the *k* nearest nodes by spatial distance.

For span masking, we randomly select continuous spans of up to 30 amino acids until 15% of input backbone coordinates are masked. Such a span masking scheme has shown to improve performance on natural language processing benchmarks (Joshi et al., 2020). The span lengths are sampled from a geometric distribution Geo(*p*) where *p* = 0.05 (corresponding to an average span length of 1/*p* = 20). The starting points for the spans are uniformly randomly sampled. Compared to independent random masking, span masking is better for GVP-Transformer but not for GVP-GNN (Table C.1).

For the amino acids with masked coordinates, we exclude the corresponding nodes from the input graph to the pre-processing GVP message passing layers, and then impute zeros for the geometric features when passing the GVP outputs into the main Transformer. Imputing zeros for missing vector features ensure the rotation- and translation-invariance of the model.

**Table B.1.** Fixed backbone sequence design performance on the more stringent structurally held-out test set from CATH v4.3 chains (and its short and single-chain subsets) in terms of per-residue perplexity (lower is better) and recovery (higher is better).

| Model | Data | Perplexity |  |  | Recovery % |  |  |
| --- | --- | --- | --- | --- | --- | --- | --- |
|  |  | Short | Single-chain | All | Short | Single-chain | All |
| Structured GNN | CATH | 10.08 | 7.04 | 7.06 | 27.8% | 35.1% | 35.4% |
| GVP-GNN | CATH | 8.13 | 5.76 | 5.86 | 31.5% | 41.1% | 40.4% |
|  | + AlphaFold2 | 9.87 | 6.61 | 6.50 | 26.3% | 36.3% | 36.8% |
| GVP-GNN-large | CATH | 8.87 | 6.62 | 6.68 | 31.0% | 37.2% | 37.4% |
|  | + AlphaFold2 | 7.08 | 4.46 | 4.39 | **34.1%** | 48.2% | 48.7% |
| GVP-Transformer | CATH | 8.80 | 6.78 | 6.97 | 28.5% | 36.7% | 36.3% |
|  | + AlphaFold2 | **6.99** | **4.36** | **4.34** | 33.0% | **48.9%** | **49.5%** |

### A.3. Details on model architectures

#### Autoregressive modeling

GVP-GNN and GVP-Transformer both have encoder-decoder architectures. The encoder only receives the structural features. The decoder receives the encoder output along with the one-hot encoding of the amino acids. In the autoregresive decoder, sequence information only propagate from amino acid *i* to *j* for *i < j*. The last decoder layer produces a 20-way scalar output per position and softmax activation to predict the probabilities for the amino acid identity at the next position in the sequence.

#### Invariance to rotation and translation

The input features for both GVP-GNN and GVP-Transformer are translation-invariant, making the overall models also invariant to translations.

Each GVP-GNN layer is rotation-equivariant, that is, for a vector feature *x* and any arbitrary rotation *T*, *Tf*(*x*) = *f*(*Tx*). With equivariant intermediate layers and an invariant output projection layer, GVP-GNN is overall invariant to rotations, since the composition of an equivariant function *f* with an invariant function *g* produces an invariant function *g*(*f*(*x*)).

The GVP-Transformer architecture is also invariant to rotations and translations. The initial GVP-GNN layers in GVP-Transformer output rotation-invariant scalar features and rotation-equivariant vector features for each amino acid. To make the overall GVP-Transformer invariant, we perform a change of basis on GVP-GNN vector outputs to produce rotation-invariant features for the Transformer. More specifically, for each amino acid, we define a local reference frame based on the N, CA, and C atom positions in the amino acid, following Algorithm 21 in AlphaFold2 (Jumper et al., 2021). We then perform a change of basis according to this local reference frame, rotating the vector features in GVP-GNN outputs into the local reference frames of each amino acid. (If GVP-GNN outputs are used directly as Transformer inputs without this change of basis, the GVP-Transformer model would not be rotation-invariant.) We concatenate this rotated “local version” of vector features together with the scalar features as inputs to the Transformer. The concatenated features are invariant to both translations and rotations on the input backbone coordinates, forming a *L × E* matrix where *L* is the number of amino acids in the protein backbone and *E* is the feature dimension. For amino acids with masked or missing coordinates, the features are imputed as zeros.

#### Transformer

We closely followed the original autoregressive encoder-decoder Transformer architecture (Vaswani et al., 2017) except for using learned positional embeddings instead of sinusoidal positional embeddings, attention dropout, and layer normalization inside the residual blocks (“pre-layernorm”). For model scaling experiments, we followed the model sizes in (Turc et al., 2019), and chose the 142-million-parameter model with 8 encoder layers, 8 decoder layers, 8 attention heads, and embedding dimension 512 based on the best validation set performance (Figure 6c shows test set ablation).

The GVP-GNN, GVP-GNN-large, and GVP-Transformer models used in the evaluations in this manuscript are all trained to convergence, with detailed hyperparameters listed in Table A.1.

**Figure B.1.**
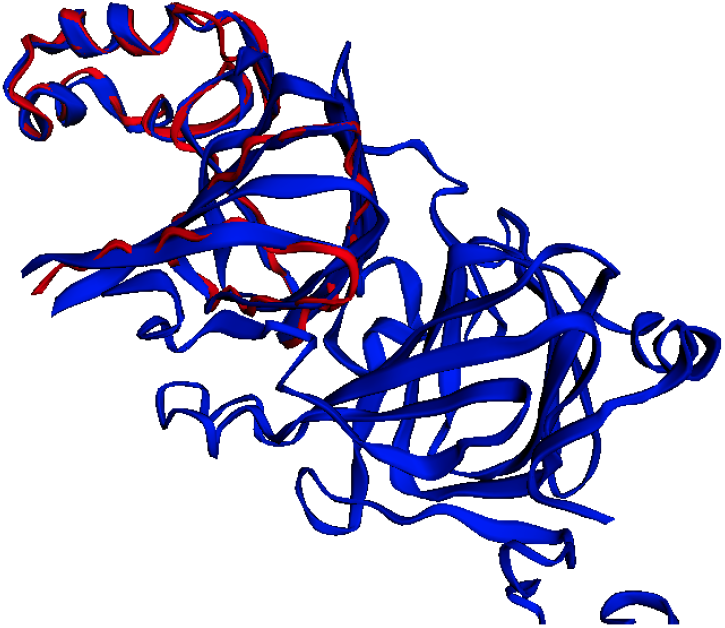
An illustrative example of structural overlap between CATH topology splits. The jack bean canavalin (PDB code 1DGW; chain Y; red) and the soybean *β*-Conglycinin (PDB code 1UIJ; chain B; blue) are assigned different topology codes in CATH (1.10.10 and 2.60.120), but they align with TM-score 0.94 and CA RMSD 0.7A on a segment of 90 residues. The difference in topology classifications likely resulted from CATH annotating only a 37-residue mainly helical segment of the jack bean canavalin as a domain while annotating a longer 176-residue mainly beta sheet segment of the soybean *β*-Conglycinin as a domain.

**Figure B.2.**
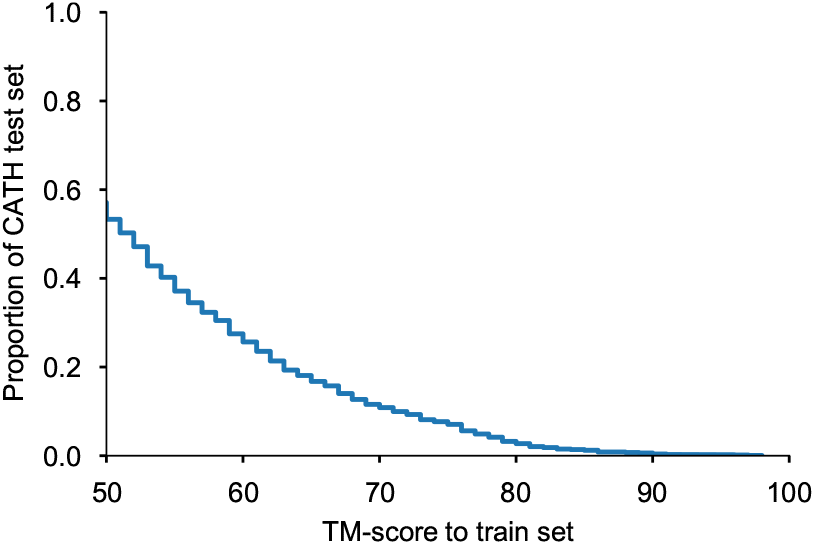
Distribution of the highest TM-score from each test example to the train set. For example, 54% of the CATH topology split test set has at least one match in the train set with TM-score above 0.5, and 27% of the topology split test set has at least one match in the train set with TM-score above 0.6.

**Table C.1.**
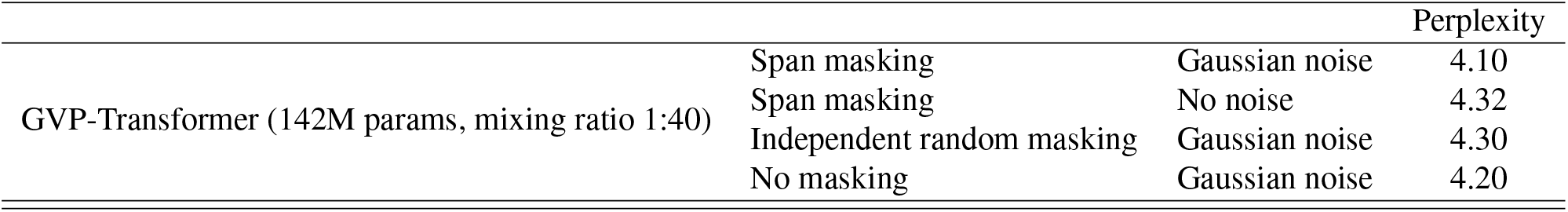
Effects of adding Gaussian noise to predicted structures and effects of span masking during training, as measured by perplexity on CATH topology split test set.

## B. TM-score-based test set

In addition to the CATH topology-based test set following previous work (Ingraham et al., 2019; Jing et al., 2021b), we also create an even more stringent test set based on pairwise TM-score comparison between train and test examples. The CATH topology split does not completely prevent high TM-score matches between train and test structures. We illustrate such an example in Figure B.1, and show overall TM-score statistics Figure B.2.

We constructed a TM-score-based test set of 223 proteins with no TMalign matches (TM-score ≥ 0.5) from the train set, using the foldseek (van Kempen et al., 2022) TMalign tool with default parameters for the pairwise search.

We found that the conclusions about model performance overall remains the same on this TM-score-based test set as on the CATH topology split test set. For consistency with prior work, we report metrics on the CATH topology test set in the main manuscript, while showing metrics on the smaller TM-score-based test set in Table B.1.

## C. Additional results and details

### Ablation on noise and masking during training

We found that GVP-Transformer models trained with Gaussian noise during training perform slightly better at test time than those trained without (Table C.1). When given full backbone coordinates at test time, training with span masking only very slightly improves model performance compared to no masking or to random masking, even though there is a much larger performance gap between random masking and span masking on regions with masked backbone coordinates (Figure 4).

### Dual-state design test set from PDBFlex

We test design performance on multiple conformations by finding test split proteins with distinct conformations in the PDBFlex database. From PDBFlex, we looks for experimental structures of protein sequences in the CATH topology split test set (95% sequence identity or above), and take all paired instances that are at least 5 angstroms apart in overall RMSD between conformations. We report perplexity on locally flexible residues (defined as local RMSD above 1 angstrom). To be more conservative in our evaluation, we show the better of the two conformations to represent single-state perplexity in Figure 7.

### Ablation on the number of GVP-GNN encoder layers in GVP-Transformer

Increasing the number of GVP-GNN encoder layers improves the overall model performance (Figure C.1), indicating that the geometric reasoning capability in GVP-GNN is complementary to the Transformer layers.

**Figure C.1.**
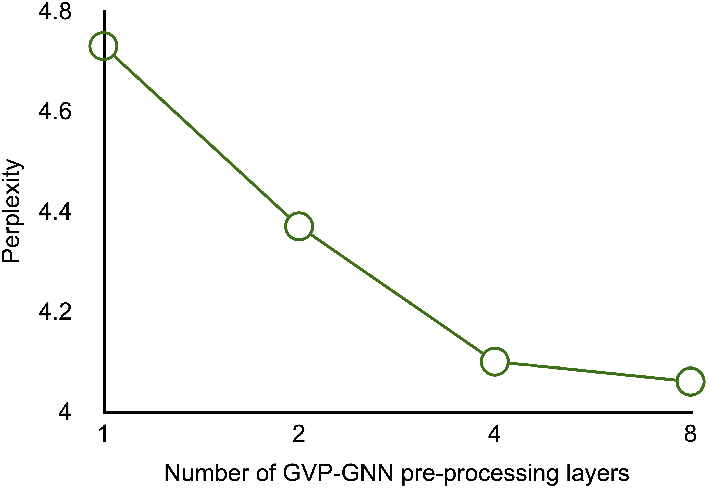
Effects of varying the number of GVP-GNN pre-processing layers in the GVP-Transformer model, as measured by perplexity on CATH topology split test set.

**Table C.2.**
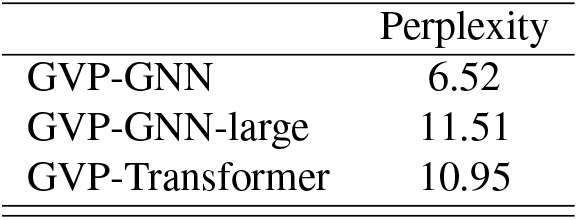
Model performance when trained only using the 12 million predicted structures without CATH training data, as measured by perplexity on CATH topology split test set.

|  | Perplexity |
| --- | --- |
| GVP-GNN | 6.52 |
| GVP-GNN-large | 11.51 |
| GVP-Transformer | 10.95 |

**Table C.3.**
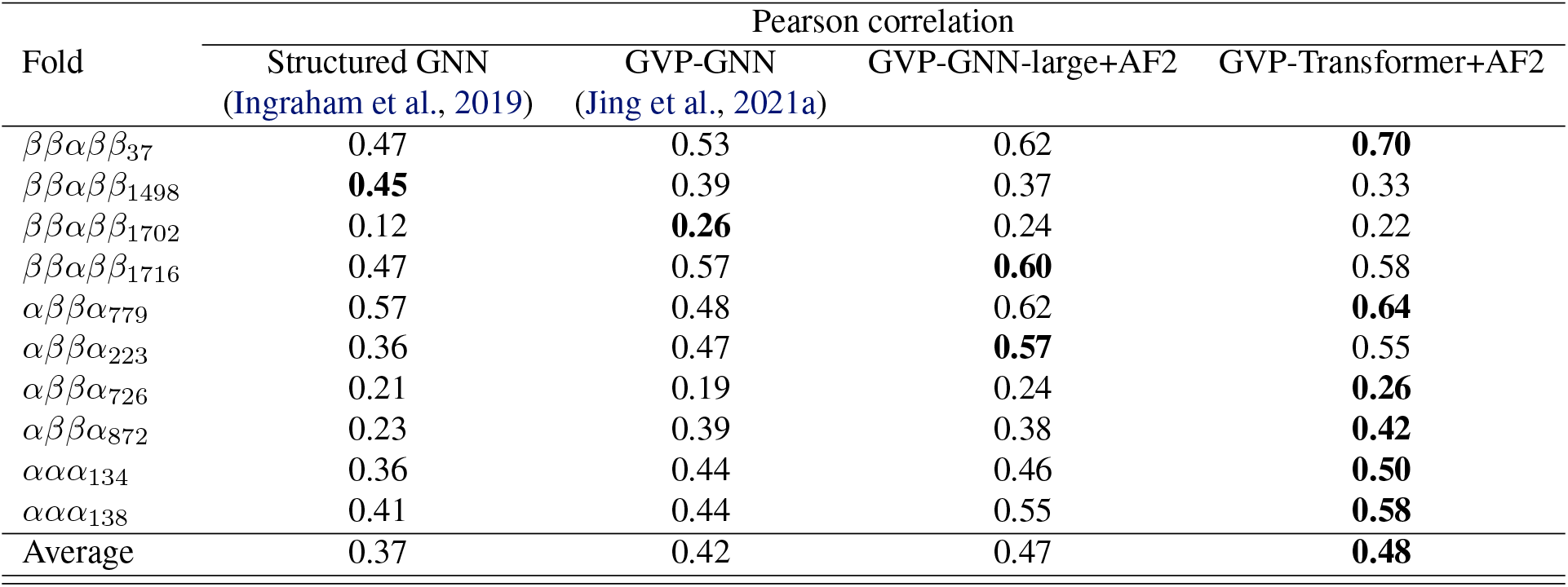
Mutation stability prediction performance for small *de novo* proteins (Rocklin et al., 2017), with highest correlation bolded.

### Model performance when trained only on predicted structures

When trained on the 12 million predicted structures without including any of the experimental structures from CATH in training data, the model performance of GVP-GNN, GVP-GNN-large, and GVP-Transformer is across the board substantially worse than when trained only on the CATH structures (Table C.2). This gap is especially pronounced for the larger GVP-GNN-large and GVP-Transformer models.

### Stability prediction on de novo small proteins

We predict protein stability on an experimentally measured stability dataset for *de novo* small proteins (Rocklin et al., 2017). We use the relative difference in sequence conditional log-likelihoods as a predictor for stability and compute Pearson correlation with the mutation effect following (Ingraham et al., 2019), assuming that more stable sequences should score higher in log-likelihoods. For each fold, Rocklin et al. (2017) starts with a reference protein and generates sequence variants with single amino acid substitutions. We calculate the Pearson correlation between sequence conditional log-likelihood scores and experimental stability measurements for all designed sequences in each fold. With predicted structures as additional training data, the GVP-Transformer model improves the pearson correlation on 8 out of the 10 folds.

### Perplexity and sequence recovery of SARS-CoV-2 RBD

We show perplexity and sequence recovery on the SARS-CoV-2 protein receptor binding domain (RBD) as an example for inverse folding. The RBD can exist in a closed-state with the RBD down or in an open-state with the RBD up (Walls et al., 2020), as illustrated in Figure C.3. The SARS-Cov-2 spike protein structure has no match with the training data with TM-score above 0.5. The SARS-Cov-2 spike protein has both an open and closed state (open state: PDB 6XRA; closed state: PDB 6VXX). We evaluate perplexity and sequence recovery conditioning on each of the two states independently and jointly. Conditioning on the open state results in better perplexity and sequence recovery than conditioning on the closed state. Conditioning on both states gives improvement in both perplexity and sequence recovery compared to conditioning only on the open state. See Table C.8 for a list of randomly sampled dual-state sequence designs from GVP-Transformer as examples.

### Predicting RBD-ACE2 binding affinity

We used the binding affinity dataset provided by Starr et al. (2020) (https://github.com/jbloomlab/SARS-CoV-2-RBD_DMS), restricting to sites within the RBM subsequence. We used the RBD-ACE2 structure determined by Lan et al. (2020) (PDB: 6M0J). For mutational effect predictions with ESM-1v, ESM-1b, and ESM-MSA-1b, we scored mutations using the masked-marginal likelihood ratio between the mutant and wildtype amino acids. To generate the MSA used as input to ESM-MSA-1b, we searched `uniclust30_2017_07` (Mirdita et al., 2017) with `hhblits` (Steinegger et al., 2019) (using two iterations and an E-value cutoff of 0.001) based on the RBD wildtype sequence as the query.

**Table C.4.** Perplexity and sequence recovery on the SARS-Cov-2 spike protein receptor binding domain (RBD), conditioned on either the closed state, the open state, or both states (illustrated in Figure C.3). The inputs to inverse folding models consist of the backbone coordinates for the entire spike protein, while the perplexity evaluation is only on the RBD.

|  | Perplexity |  |  | Recovery % |  |  |
| --- | --- | --- | --- | --- | --- | --- |
|  | Open state | Closed state | Dual-state | Open-state | Closed state | Dual-state |
| GVP-GNN | 4.64 | 5.13 | 4.20 | 45.3% | 44.2% | 49.7% |
| GVP-Transformer | 4.50 | 4.96 | 4.06 | 49.2% | 48.1% | 53.6% |

**Table C.5.** Protein complex stability on SKEMPI test set (binary classification of increase in stability on single-point mutations). Although only trained on single chains, the inverse-folding models generalize to protein complexes. Giving the full complex as input, *complex*, improves performance compared to giving only the chain as input, *chain*. Zero-shot prediction compared to fully supervised and supervised transfer learning methods from (Townshend et al., 2020) and (Jing et al., 2021a) trained on the SKEMPI train set.

|  |  | AUROC |
| --- | --- | --- |
| Supervised | 3DCNN | 0.57 |
|  | GNN | 0.62 |
|  | ENN | 0.57 |
|  | GVP-GNN | 0.68 |
| Transfer | GVP-GNN | **0.71** |
| Zero-shot | GVP-GNN (chain) | 0.58 |
|  | GVP-GNN (complex) | **0.71** |
|  | GVP-GNN-large+AF2 (chain) | 0.61 |
|  | GVP-GNN-large+AF2 (complex) | **0.71** |
|  | GVP-Transformer+AF2 (chain) | 0.60 |
|  | GVP-Transformer+AF2 (complex) | 0.68 |

**Table C.6.** Zero-shot performance on AAV split (Dallago et al., 2021).

| Evaluation subset | Spearman correlation (zero-shot) |  |  |  |
| --- | --- | --- | --- | --- |
|  | ESM-1v | GVP-GNN | GVP-GNN-large+AF2 | GVP-Transformer+AF2 |
| Mutated | $-0.23 \pm 0.03$ | **$0.34 \pm 0.02$** | $0.29 \pm 0.03$ | **$0.31 \pm 0.03$** |
| Designed | $0.42 \pm 0.02$ | $0.65 \pm 0.01$ | **$0.72 \pm 0.01$** | $0.67 \pm 0.02$ |
| High-fitness | $0.22 \pm 0.02$ | $0.13 \pm 0.03$ | $0.21 \pm 0.03$ | **$0.26 \pm 0.02$** |
| Sampled | $-0.21 \pm 0.03$ | **$0.35 \pm 0.02$** | $0.30 \pm 0.02$ | $0.30 \pm 0.03$ |
| $\geq 2$ mutations | $-0.20 \pm 0.03$ | **$0.35 \pm 0.03$** | $0.29 \pm 0.04$ | $0.30 \pm 0.02$ |
| $\geq 3$ mutations | $0.28 \pm 0.03$ | $0.53 \pm 0.02$ | **$0.62 \pm 0.02$** | **$0.64 \pm 0.02$** |
| $\geq 8$ mutations | $0.20 \pm 0.03$ | $0.47 \pm 0.02$ | **$0.53 \pm 0.02$** | **$0.55 \pm 0.02$** |

**Figure C.2.**
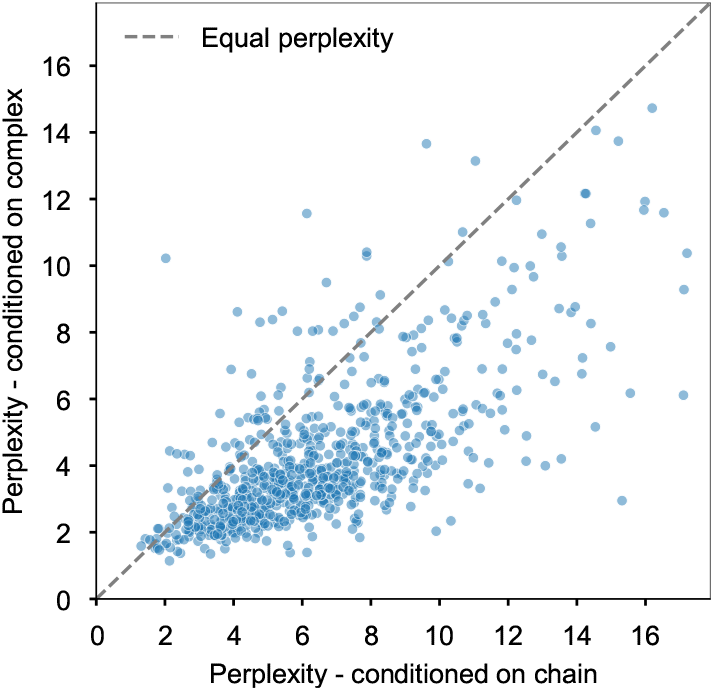
Fixed backbone sequence design perplexity for protein complexes. The model is evaluated on 796 structurally held-out protein complexes. Comparison of conditioning on the backbone coordinates of individual chains (x-axis) with conditioning on backbone coordinates of the entire complex (y-axis). Note that for both values perplexity is evaluated on the same chain in the complex. The shift to the lower right indicates improved perplexity when the model is given the complete structure of the complex.

**Figure C.3.**
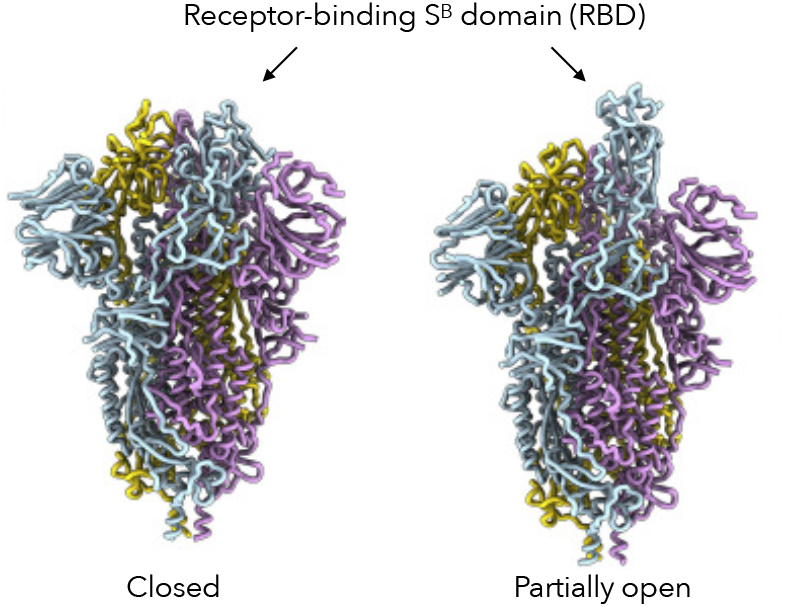
Illustration of the closed and open states of the SARS-CoV-2 spike protein receptor-binding domain. Cryo-EM structures from (Walls et al., 2020) (open state: PDB 6XRA; closed state: PDB 6VXX).

### Predicting complex stability changes upon mutations

SKEMPI (Jankauskaiteė et al., 2019) is a database of binding free energy changes upon single point mutations within protein complex interfaces. This database is used as a task in the Atom3D benchmark suite (Townshend et al., 2020) for comparing supervised stability prediction methods. The task is to classify whether the stability of the complex increases as a result of the mutation. We compare zero-shot predictions using inverse folding models to supervised and transfer learning methods (Townshend et al., 2020; Jing et al., 2021a) on the Atom3D test set. We find that sequence log-likelihoods from GVP-GNN, GVP-GNN-large, and GVP-Transformer models are all effective zero-shot predictors of stability changes of protein complexes (Table C.5), performing comparably to the best supervised method which uses transfer learning.

### Predicting insertion effects on AAV

Using masked coordinate tokens at insertion regions, inverse folding models can also predict the effects of sequence insertions. Adeno-associated virus (AAV) capsids are a promising gene delivery vehicle, approved by the US Food and Drug Administration for use as gene delivery vectors in humans. Focusing on mutating a 28-amino acid segment, Bryant et al. (2021) generated more than 200,000 variants of AAV sequences with 12–29 mutations across this region, and measured their ability to package of a DNA payload. This dataset is unique compared to many other mutagenesis datasets in that most sequences feature random insertions in the 28-amino acid segment, as opposed to only random substitutions.

We use inverse folding models to predict insertion and substitution effects as follows: For each sequence, we input the full backbone coordinates of the wild-type (PDB: 1LP3), and insert one masked token into the input backbone coordinates for each insertion. Then we compare the conditional sequence log-likelihood on this input with masks to the conditional sequence log-likelihood of the wild-type sequence on the wild-type backbone. The difference in these two conditional log-likelihoods are used as the score for predicting packaging ability.

We report the zero-shot performance on each of the 7 data subsets evaluated in the FLIP (Dallago et al., 2021) benchmark suite. For amino acid insertions (marked as lowercase letters in the FLIP data), the corresponding backbone coordinates for those amino acids are marked as unknown in the input structure. As shown in Table C.6, GVP-Transformer trained with predicted structures outperforms the sequence-only zero-shot prediction baseline ESM-1v on 6 out of the 7 data subsets. The reported standard deviations are calculated by sampling different subsets of 10,000 variants from the evaluation data.

For ESM-1v, we scored variant sequences based on the independent marginals formula as described in Equation 1 from Meier et al. (2021), scoring mutations using the log odds ratio at the mutated position, assuming an additive model when a set of multiple mutations *T* exist in the same sequence:

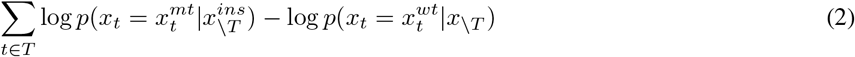

where 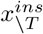 in the first term is the wild-type sequence with mask tokens at insertion positions and *x*_\T_ in the second term is the wild-type sequence without insertions.

### Confusion matrix

We calculated the substitution scores between native sequences and sampled sequences (sampled with temperature *T* = 1) by using the same log odds ratio formula as in the BLOSUM62 substition matrix. For two amino acids *x* and *y*, the substitution score *s*(*x, y*) is

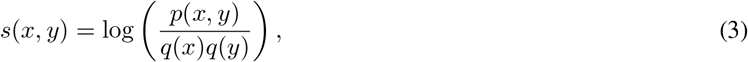

where *p*(*x, y*) is the jointly likelihood that native amino acid *x* is substituted by sampled amino acid *y*, *q*(*x*) is the marginal likelihood in the native distribution, and *q*(*y*) is the marginal likelihood in the sampled distribution.

### Calibration

Calibration curves examines how well the probabilistic predictions of a classifier are calibrated, plotting the true frequency of the label against its predicted probability. When computing the calibration curve, for each amino acid, we bin the predicted probabilities into 10 bins and then compare with the true probability.

### Placement of hydrophobic residues

We define the amino acids IVLFCMA as hydrophobic residues, and inspect the distribution of solvent accessible surface area for both hydrophobic residues and polar (non-hydrophobic) residues. Solvent accessible surface area calculated with the Shrake-Rupley (“rolling probe”) algorithm from the biotite package (Kunzmann & Hamacher, 2018) and summed over all atoms in each amino acid. All models have similar distributions of accessible surface area for hydrophobic residues, also similar to the distribution in native sequences (Figure C.6).

### Sampling speed

We profile the sampling speed with PyTorch Profiler, averaging over the sampling time for 30 sequences in each sequence length bucket on a Quadro RTX 8000 GPU with 48GB memory. For the generic Transformer decoder, we use the incremental causal decoding implementation in fairseq (Ott et al., 2019). For GVP-GNN, we use the implementation from the gvp-pytorch GitHub repository.

**Figure C.4.**
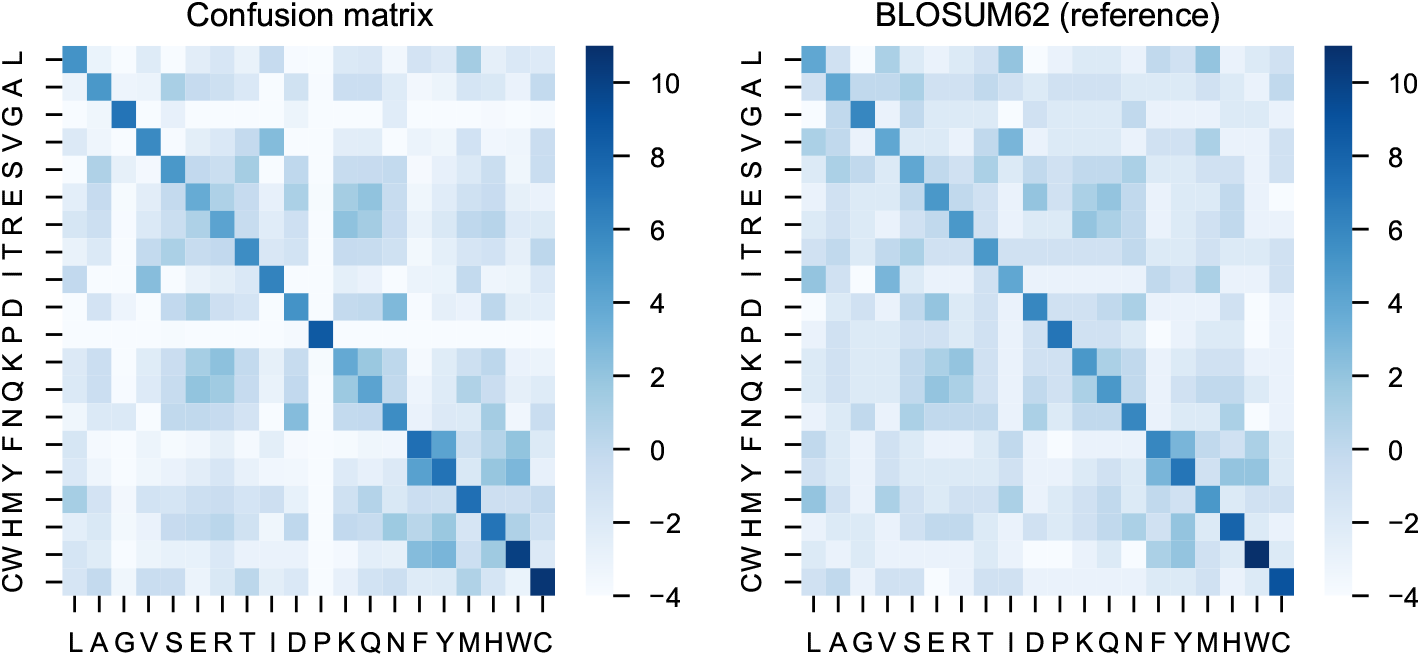
Confusion matrix between native sequence and sampled sequences from the model, compared to BLOSUM62 as reference.

**Figure C.5.**
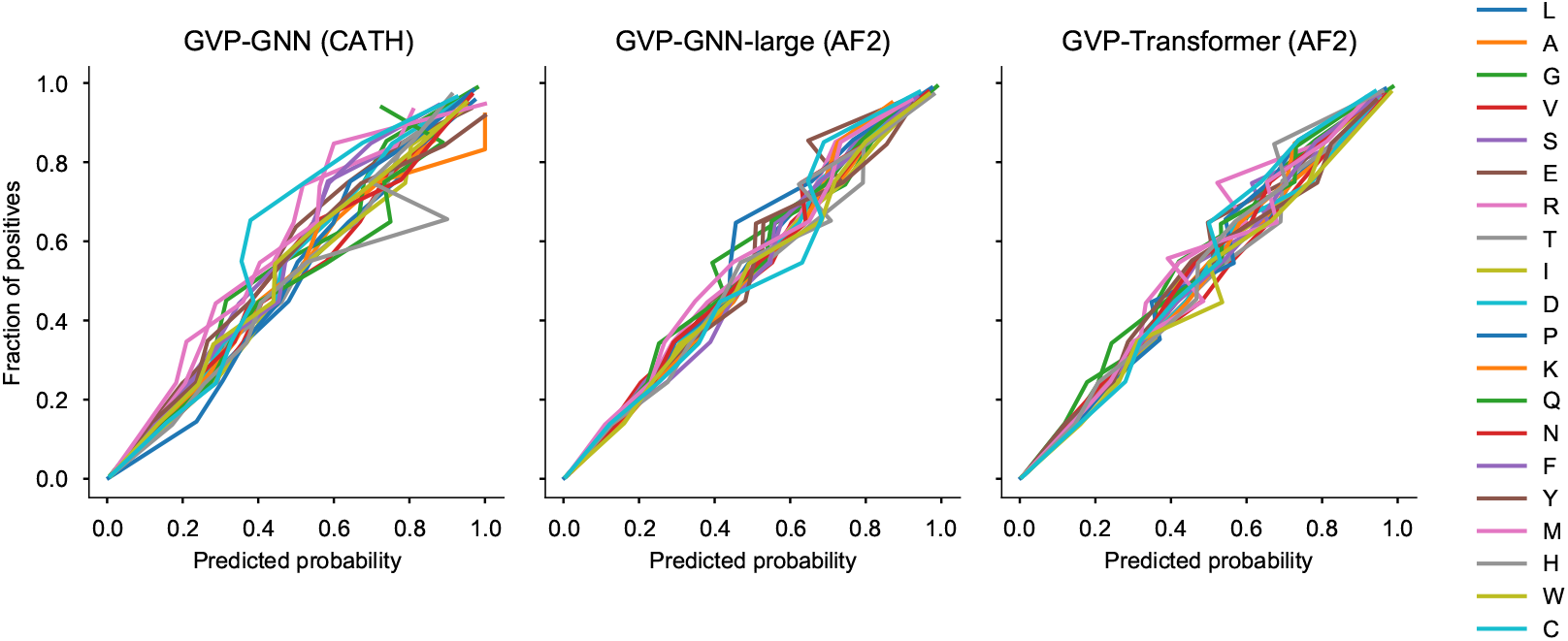
Calibration.

**Figure C.6.**
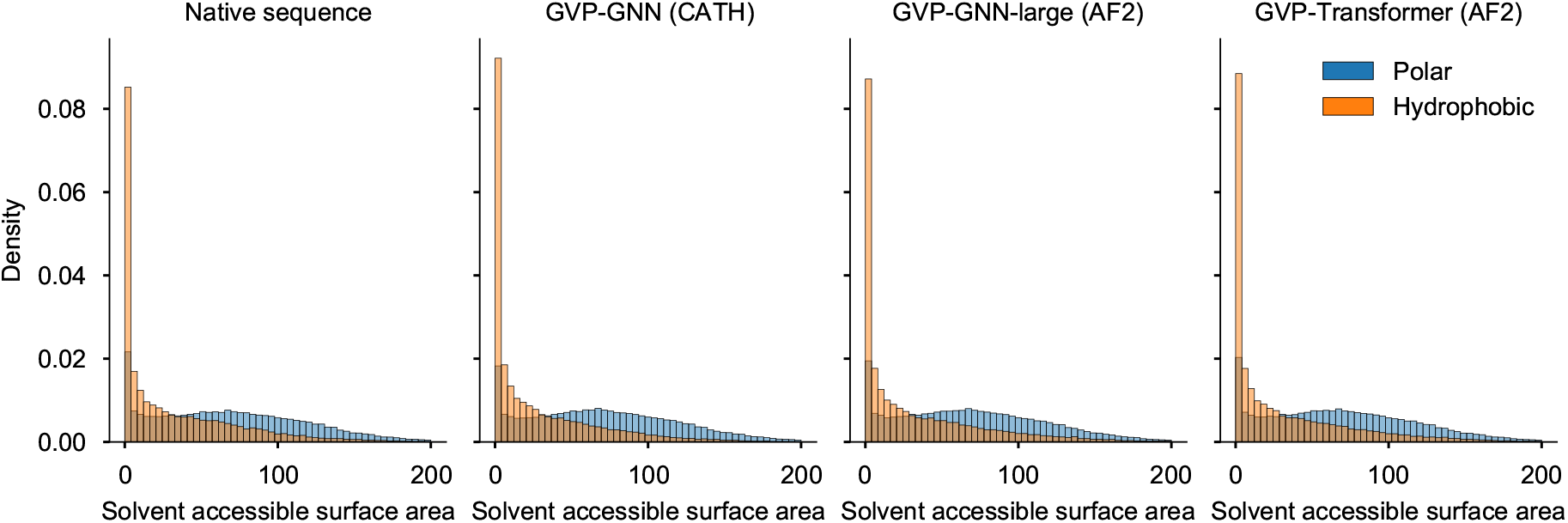
The majority of hydrophobic residues are buried, following a long tail accessible surface area distribution as in native sequences.

**Table C.7.** Average time required for sampling one sequence, using open source implementation of GVP-GNN and open source implementation of Transformer from fairseq (Ott et al., 2019).

| Sequence length | Average sampling time per sequence |  |  |  |  |
| --- | --- | --- | --- | --- | --- |
| | $\leq 100$ | 100 – 200 | 200 – 300 | 300 – 400 | 400 – 500 |
| GVP-GNN (3 layers) | 3.7s | 9.3s | 20.8s | 76.9s | 150.3s |
| GVP-GNN-large (8 layers) | 6.7s | 11.8s | 47.5s | 90.3s | 168.8s |
| GVP-Transformer (8 layers) | 1.5s | 2.6s | 9.0s | 16.2s | 26.0s |

**Table C.8.**
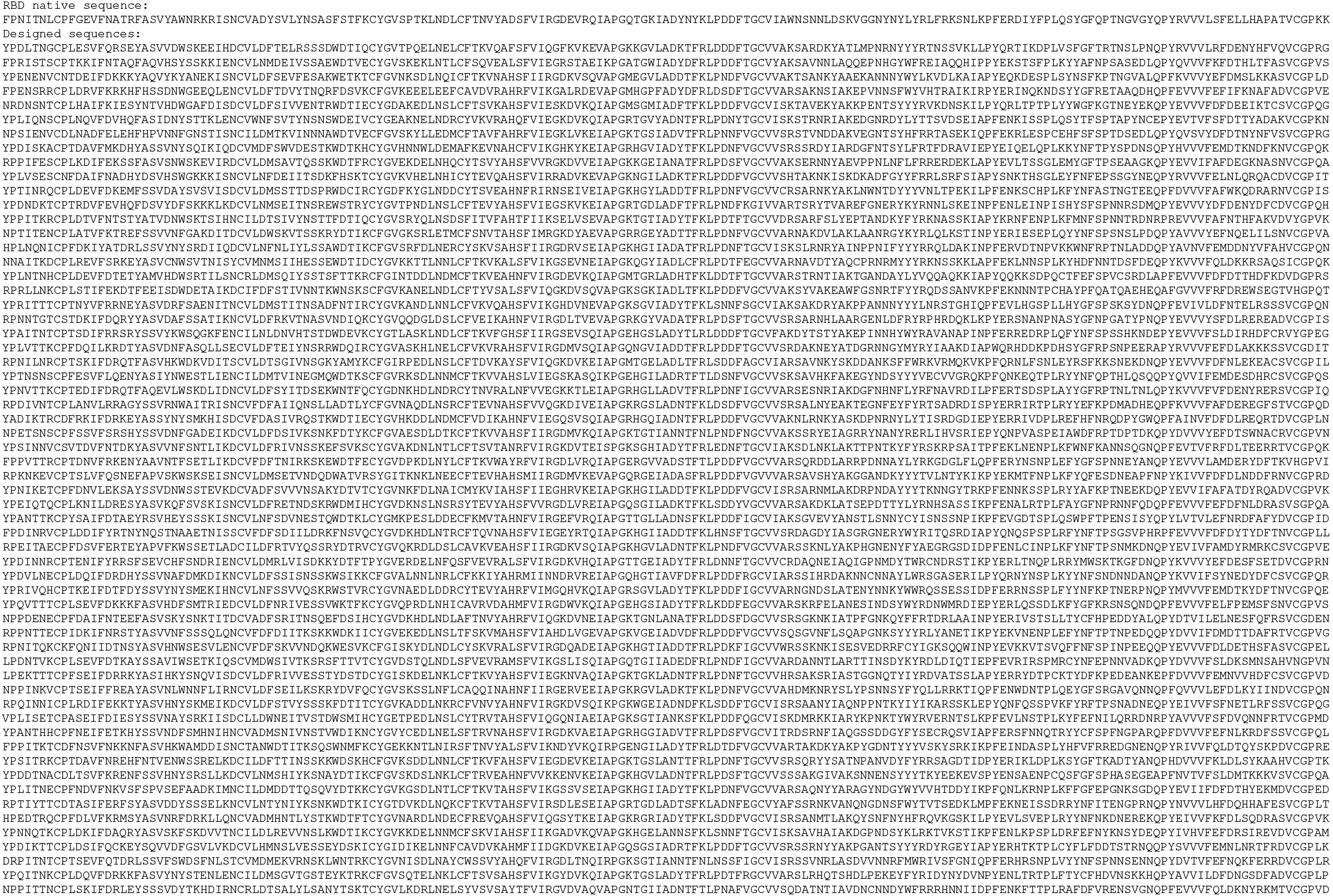
60 randomly sampled RBD dual-state sequence designs from the GVP-Transformer model with sampling temperature 1.0 and conditioned on both the open and closed states.

